# Additive effects on craniofacial development upon conditional ablation of PDGFR*α* and SHP2 in the mouse neural crest lineage

**DOI:** 10.1101/2025.02.13.638176

**Authors:** Daniel Fuhr, Jessica Johnston, Elliott P. Brooks, Katherine A. Fantauzzo

## Abstract

**Background:** Activity of the receptor tyrosine kinase PDGFR*α* and the tyrosine phosphatase SHP2 are critical for vertebrate craniofacial development. We sought to determine the effect of SHP2 binding to PDGFR*α* via phenotypic and biochemical analyses of an allelic series of mouse embryos with combined loss of both proteins in the neural crest lineage.

**Results:** We demonstrated that SHP2 preferentially binds PDGFR*α*/*α* homodimers among the three PDGFR dimers. Analysis of allelic series mutant embryos revealed increased cell death in the lateral nasal and maxillary processes at E10.5, variably penetrant facial blebbing, facial hemorrhaging, midline clefting and loss of the mandibular region at E13.5, and widespread craniofacial bone and cartilage defects at birth. Further, we showed that loss of SHP2 leads to increased phosphorylation of PDGFR*α* and the downstream effector Erk1/2 in E10.5 allelic series mutant embryo lysates.

**Conclusions:** Together, our findings demonstrate additive effects on craniofacial development upon conditional ablation of PDGFR*α* and SHP2 in the mouse neural crest lineage and indicate that SHP2 may negatively and positively regulate PDGFR*α* signaling through distinct mechanisms.

## Introduction

The mammalian platelet-derived growth factor (PDGF) family contains two receptor tyrosine kinases (RTKs), PDGFR*α* and PDGFR*β*, each of which possesses an extracellular ligand-binding domain, a transmembrane domain and an intracellular domain^1^. The intracellular portion of the PDGFRs contains split tyrosine kinase domains that become activated upon ligand binding and receptor dimerization, leading to the autophosphorylation of intracellular tyrosine residues. Once phosphorylated, these tyrosine residues are bound by signaling molecules that, in the case of PDGFR*α*, include Src family tyrosine kinases, the tyrosine phosphatase SHP2, phosphatidylinositol 3-kinase (PI3K), the adaptor protein Crk and phospholipase C*γ*^2^. Heterozygous missense variants in the coding region of *PDGFRA* as well as single base-pair substitutions in the 3’ untranslated region have been associated with nonsyndromic cleft palate in humans^3^. Similarly, conditional ablation of *Pdgfra* in the murine neural crest cell (NCC) lineage, which includes the craniofacial mesenchyme, using the *Wnt1- Cre^Tg^* allele^4^ leads to facial clefting, facial hemorrhaging, thymus hypoplasia and aortic arch defects^5,6^. Studies examining the function of PDGFR*α* in the NCC lineage have revealed roles in promoting migration of cranial NCCs, proliferation and survival of the craniofacial mesenchyme and osteoblast differentiation^6–10^. The interaction between PDGFR*α* and a subset of associated signaling molecules has been explored in the context of craniofacial development, particularly in the case of PI3K^7,11–13^. However, the effect of SHP2 binding to PDGFRs, especially PDGFR*α*, is unclear, as SHP2 has been shown to both positively and negatively regulate PDGFR signaling. SHP2 binds human PDGFR*α* at Y720 and Y754 and human PDGFR*β* at Y1009 (Y1008 in mice)^14–17^. Upon binding to PDGFR*α* and PDGFR*β* in various cell culture contexts, SHP2 can act as an adapter protein to recruit Grb2, resulting in the activation of Ras in the case of PDGFR*β*^14,18–20^. Alternatively, SHP2 binding to PDGFR*β*also results in dephosphorylation of the autophosphorylated receptor in *in vitro* kinase assays and upon PDGF-BB ligand stimulation of HepG2 human hepatocellular carcinoma cells^21^.

Balanced activity of the ubiquitously expressed, non-transmembrane tyrosine phosphatase SHP2^22^ is critical for proper craniofacial development in both humans and mice, as evidenced by phenotypes resulting from either loss- or gain-of-function variants in *PTPN11*, the gene encoding SHP2. In humans, variants in *PTPN11* that impair its protein tyrosine phosphatase catalytic activity result in LEOPARD syndrome-1 (LPRD1; OMIM 151100)^23^, which is characterized by a wide range of manifestations that include ocular hypertelorism and sensorineural deafness^24^. Modeling of the two most common of these LPRD1-associated variants in mice results in an increased skull width-to-length ratio, small, slanted eyes, increased interorbital distance and a planar nasal bridge, among other defects^25,26^. Moreover, conditional ablation of *Ptpn11* in the murine NCC lineage using the *Wnt1-Cre* driver leads to embryonic lethality after mid-gestation and severely hypoplastic or absent anterior craniofacial bones and cartilage^27,28^. *Sox10* RNA *in situ* hybridization and Cre reporter-based lineage tracing analyses of control versus conditional homozygous mutant embryos at E9.5-E10.0 revealed no differences in the extent of NCC migration into the frontonasal prominence nor pharyngeal arches^27,28^, and immunofluorescence analyses at E9.5 demonstrated no differences in the levels of Ki67 nor cleaved caspase 3 in the pharyngeal arches^27^, arguing against early defects in NCC specification, migration, proliferation and survival. However, phospho-Erk1/2 levels were significantly decreased in the pharyngeal arches of E9.5 embryos with conditional ablation of *Ptpn11* in the NCC lineage^27^. Conversely, gain-of-function variants in *PTPN11* underlie Noonan syndrome-1 (NS1; OMIM 163950) in humans^29^, which is characterized by dysmorphic facial features such as a broad forehead, hypertelorism, downslanting palpebral fissures, low-set, posteriorly-rotated ears and a high-arched palate, among other abnormalities^30^. Modeling in mice of a NS1-associated gain-of-function variant in *Ptpn11* results in animals with a shorter skull, a greater inner canthal distance and a wider snout, as well as increased phospho-Erk1/2 expression in the facial region at E11.5^31^. Conditional expression of a second NS1-associated variant in *Ptpn11* in the murine NCC lineage using the *Wnt1-Cre* driver leads to similar craniofacial phenotypes and hyperactivation of Erk1/2 in the frontal bones, both of which are rescued upon injection of the MEK1/2 inhibitor U0126^32^.

Here, we sought to determine the effect of SHP2 binding to PDGFR*α* via phenotypic and biochemical analyses of an allelic series of embryos with combined loss of both proteins in the NCC lineage. We demonstrated that these ablations are additive in nature, as double-homozygous mutant embryos exhibited a combination, but not an improvement or worsening, of the previously reported phenotypes observed upon conditional ablation of PDGFR*α* or SHP2 in this context. We provided multiple lines of evidence that SHP2 may negatively and positively regulate PDGFR*α*signaling. A model is put forth to explain these seemingly contradictory results in which SHP2 binds and dephosphorylates PDGFR*α* while simultaneously increasing survival through conserved Erk1/2-independent mechanisms.

## Results

We previously demonstrated that SHP2 is differentially engaged with PDGFR*α* in response to stimulation of mouse embryonic fibroblasts with various PDGF ligands^33^. Specifically, following immunoprecipitation of PDGFR*α*, we found the highest levels of bound SHP2 via western blotting following treatment with PDGF-AA, with reduced levels of SHP2 upon treatment with PDGF-BB and PDGF-DD^33^. As we demonstrated that PDGF-AA stimulation results in autophosphorylation of PDGFR*α*exclusively in this context, while PDGF-BB and PDGF-DD treatments led to the autophosphorylation of both receptors with a preference for PDGFR*β*^33^, these results imply increased binding of SHP2 upon PDGFR*α*/*α* homodimer versus PDGFR*α*/*β* heterodimer formation. Interestingly, we were never able to detect SHP2 binding to PDGFR*β* upon stimulation with the above PDGF ligands in mouse embryonic fibroblasts^33^, indicating that SHP2 may primarily exert its function on PDGFR signaling in this setting through binding to PDGFR*α*. To definitively examine SHP2 binding to the various PDGFR dimers, we employed a bimolecular fluorescence complementation (BiFC) approach that we have previously used to reveal differences in the timing and extent of PDGFR*α*/*α* homodimer, PDGFR*α*/*β* heterodimer and PDGFR*β*/*β* homodimer dimerization, autophosphorylation, trafficking and activation of downstream signaling^34,35^. This technique utilizes a split Venus fluorescent protein^36^ fused to the C-terminus of individual PDGFRs and circumvents limitations with antibody-based approaches that cannot distinguish between monomeric versus dimerized receptors, nor PDGFR homodimers versus heterodimers. Here, we transfected HEK 293T/17 cells with pairs of plasmids expressing C-terminal fusions of PDGFRs with BiFC fragments corresponding to the N-terminal (V1) or C-terminal (V2) regions of Venus, or a control plasmid expressing a membrane-targeted Venus protein with an N-terminal myristoylation tag^37^. Given that PDGF-AA ligand treatment leads to the highest levels of PDGFR*α*/*α*homodimer formation, while PDGF-BB stimulation induces the greatest extent of PDGFR*α*/*β* heterodimer and PDGFR*β*/*β*homodimer formation across multiple cellular contexts^33,35,38^, we stimulated cells expressing PDGFR*α*V1/*α*V2 with PDGF-AA and cells expressing PDGFR*α*V1/*β*V2 and PDGFR*β*V1/*β*V2 with PDGF-BB. Subsequent immunoprecipitation with a GFP-Trap nanobody that binds the recombined V1/V2 interface^36^ followed by western blotting with an anti-SHP2 antibody revealed the strongest band intensities upon PDGFR*α*/*α*homodimer formation (3.677 ± 0.6740 relative SHP2 binding, mean ± s.e.m.), reduced band intensities upon PDGFR*α*/*β* heterodimer formation (2.209 ± 0.1431) and background band intensities upon PDGFR*β*/*β*homodimer formation (0.9545 ± 0.2184) (**Figure 1A,B**). These results conclusively demonstrated that SHP2 preferentially binds PDGFR*α*/*α* homodimers among the three PDGFR dimers.

**Figure 1.**
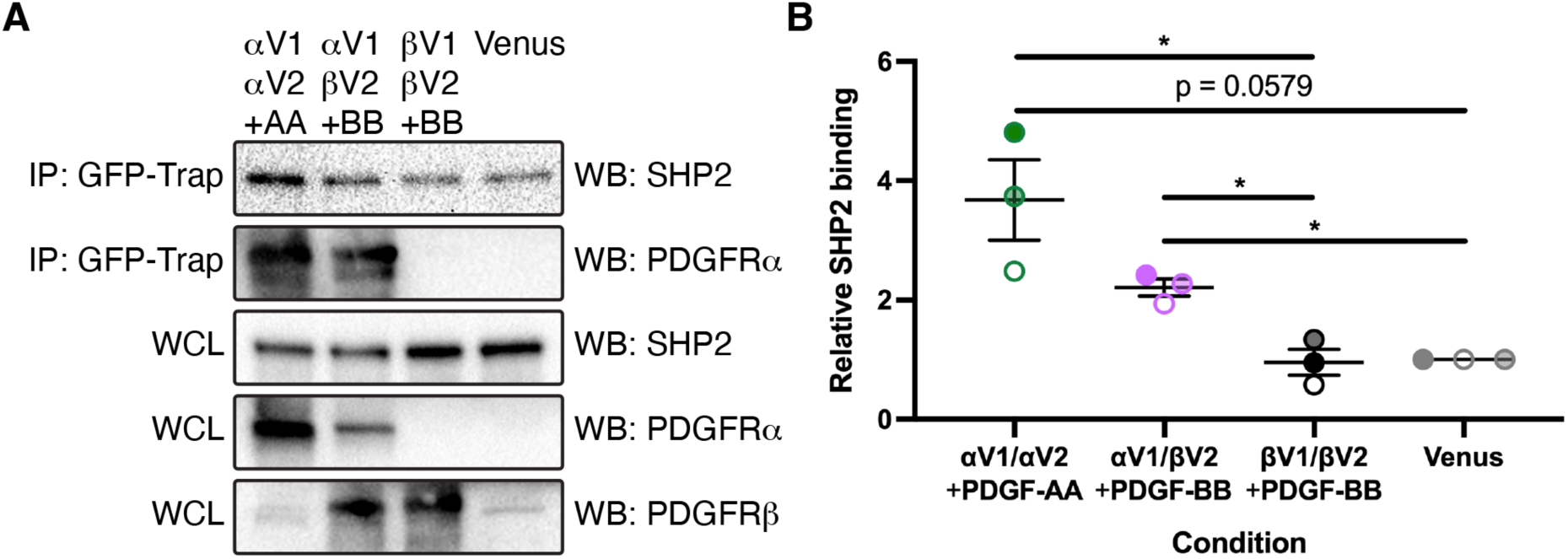
SHP2 preferentially binds PDGFR*α*/*α*homodimers. (A) Immunoprecipitation (IP) of dimerized PDGFRs or myr-Venus from whole-cell lysates (WCL) with a GFP-Trap nanobody followed by western blotting (WB) with anti-SHP2 and anti-PDGFR*α*antibodies. (B) Scatter dot plot depicting quantification of band intensities from three independent experiments as in A. Data are presented as mean ± s.e.m. *, p < 0.05. Shades correspond to independent experiments. *n* = 3 biological replicates per condition.

To ascertain the effect of SHP2 binding to PDGFR*α* in the craniofacial mesenchyme, we performed genetic epistasis experiments by intercrossing *Pdgfra^fl/fl^;Shp2^fl/fl^* mice with *Pdgfra^+/fl^;Shp2^+/fl^;Wnt1-Cre^+/Tg^* mice and analyzing the resulting progeny at E10.5, just before the onset of facial clefting phenotypes in embryos with conditional ablation of *Pdgfra* in the NCC lineage^5,6^. We found that control *Pdgfra^+/fl^;Shp2^+/fl^;Wnt1-Cre^+/+^* embryos (48 embryos vs. 28 expected embryos out of 226 total, *χ*^2^ two-tailed *p* < 0.0001), *Pdgfra^+/fl^;Shp2^+/fl^;Wnt1-Cre^+/Tg^* double-heterozygous embryos (54 embryos vs. 28 expected embryos, *χ*^2^ two-tailed *p* < 0.0001) and *Pdgfra^fl/fl^;Shp2^fl/fl^;Wnt1-Cre^+/Tg^* double-homozygous mutant embryos (38 embryos vs. 28 expected embryos, *χ*^2^ two-tailed *p* = 0.050) were significantly overrepresented among the harvested embryos, while *Pdgfra^+/fl^;Shp2^fl/fl^;Wnt1-Cre^+/Tg^* embryos were significantly underrepresented (16 embryos vs. 28 expected embryos, *χ*^2^ two-tailed *p* = 0.014; **Table 1**). Interestingly, *Pdgfra^+/fl^;Shp2^fl/fl^;Wnt1-Cre^+/+^* embryos (8 embryos vs. 28 expected embryos, *χ*^2^ two-tailed *p* < 0.0001) and *Pdgfra^fl/fl^;Shp2^+/fl^;Wnt1-Cre^+/+^* embryos (7 embryos vs. 28 expected embryos, *χ*^2^ two-tailed *p* < 0.0001) were also significantly underrepresented at this timepoint (**Table 1**). *Pdgfra^fl/fl^;Shp2^+/fl^;Wnt1-Cre^+/Tg^* embryos were recovered at Mendelian frequencies at E10.5 (**Table 1**). With the exception of double-homozygous mutant embryos, the majority of embryos across each of the remaining seven allele combinations had no gross morphological phenotype at this timepoint (**Table 1**). At E10.5, double-homozygous mutant embryos exhibited hypoplastic facial processes and narrower heads than control embryos (**Figure 2A-B’**), as well as incompletely penetrant blebbing of the surface ectoderm in the facial region (17%, n = 12), facial hemorrhaging (17%, n = 12) and loss of the mandibular processes (25%, n = 12) (**Table 1**). However, the normalized distance between nasal pits was not significantly different between control and any of the experimental genotypes containing the *Wnt1-Cre* transgene at this timepoint (**Figure 2C**), arguing against an early defect at the midline.

**Table 1.** Genotypes and phenotypes of E10.5 embryos from intercrosses of *Pdgfra^fl/fl^;Shp2^fl/fl^* mice with *Pdgfra^+/fl^;Shp2^+/fl^;Wnt1-Cre^+/Tg^* mice.

| Genotype | Expected | Observed | Normal of alive | Dead | Abnormal head | Facial bleb | Facial hemorrhage | Absent MdP |
| --- | --- | --- | --- | --- | --- | --- | --- | --- |
| $\alpha^{+/fl};S^{+/fl};W1C^{+/+}$ | 0.125 | 0.212* | 23/25 | 0 | 1/25 | 2/25 | 0/25 | 0/25 |
| $\alpha^{+/fl};S^{+/fl};W1C^{+/Tg}$ | 0.125 | 0.239* | 9/11 | 0 | 2/11 | 1/11 | 0/11 | 0/11 |
| $\alpha^{+/fl};S^{fl/fl};W1C^{+/+}$ | 0.125 | 0.035* | 3/3 | 0 | 0/3 | 0/3 | 0/3 | 0/3 |
| $\alpha^{+/fl};S^{fl/fl};W1C^{+/Tg}$ | 0.125 | 0.071* | 5/8 | 0 | 3/8 | 1/8 | 1/8 | 1/8 |
| $\alpha^{fl/fl};S^{+/fl};W1C^{+/+}$ | 0.125 | 0.031* | 2/2 | 0 | 0/2 | 0/2 | 0/2 | 0/2 |
| $\alpha^{fl/fl};S^{+/fl};W1C^{+/Tg}$ | 0.125 | 0.093 | 5/9 | 0 | 4/9 | 2/9 | 0/9 | 0/9 |
| $\alpha^{fl/fl};S^{fl/fl};W1C^{+/+}$ | 0.125 | 0.150 | 12/13 | 0 | 1/13 | 0/13 | 1/13 | 0/13 |
| $\alpha^{fl/fl};S^{fl/fl};W1C^{+/Tg}$ | 0.125 | 0.168* | 5/12 | 0 | 6/12 | 2/12 | 2/12 | 3/12 |

\* = statistically significant in $\chi^2$ analysis; n = 226 embryos harvested, n = 83 embryos scored for phenotype

**Figure 2.**
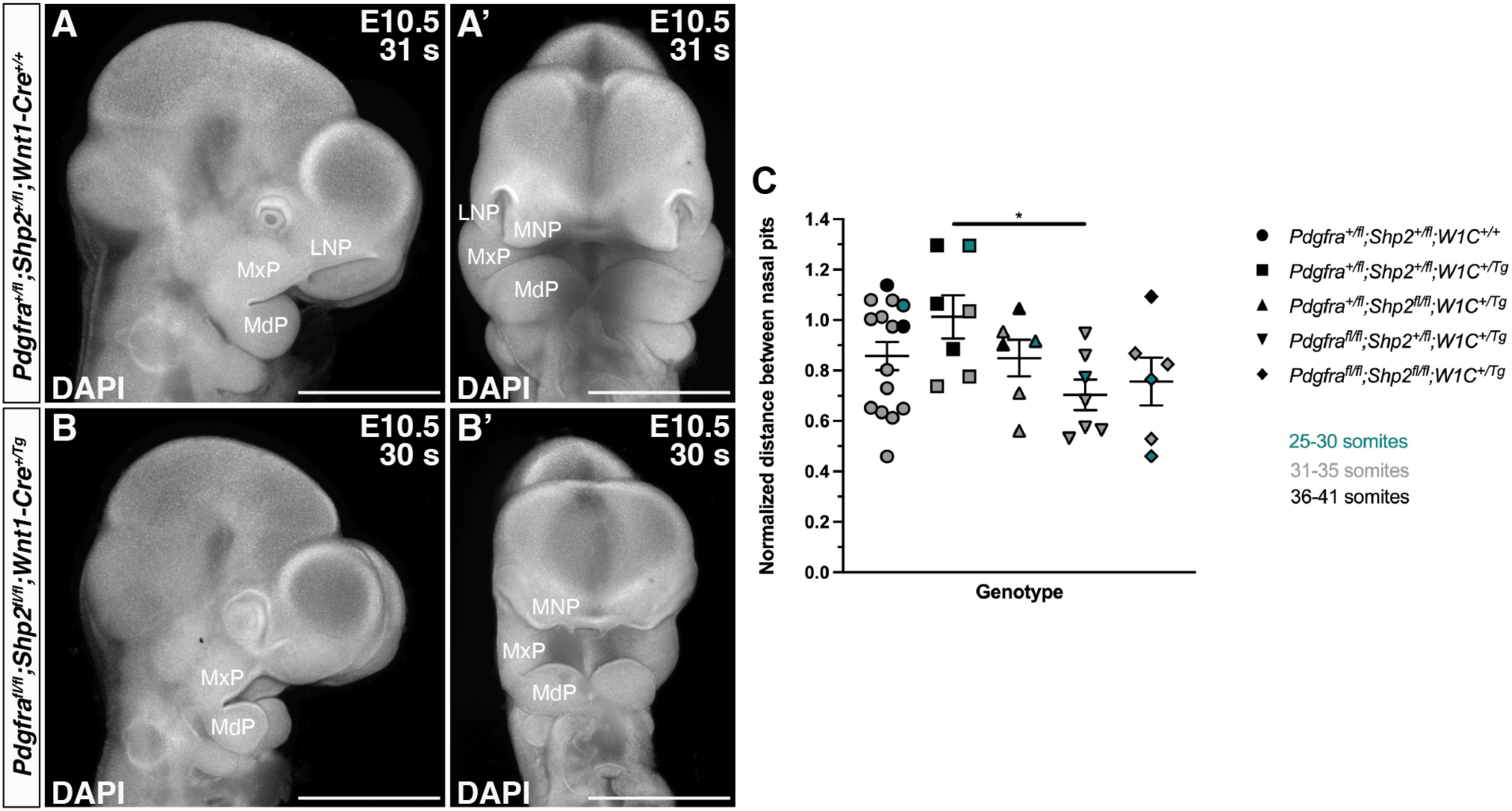
Double-homozygous mutant embryos exhibit hypoplastic facial processes and narrow heads at E10.5 with no defect at the midline. (A-B’) Gross morphology of E10.5 *Pdgfra^+/fl;^Shp2^+/fl^;Wnt1-Cre^+/+^* control embryos (A,A’) and double-homozygous mutant embryos (B,B’) viewed laterally (A,B) and frontally (A’,B’) after DAPI staining (white). s, somite pairs; LNP, lateral nasal process; MxP, maxillary process; MdP, mandibular process; MNP, medial nasal process. Scale bars, 1 mm. (C) Scatter dot plot depicting normalized distance between nasal pits across five genotypes at E10.5. Data are presented as mean ± s.e.m. *, p < 0.05. Colors correspond to number of somite pairs in assayed embryos. *n ≥* 6 biological replicates per genotype.

We next assessed the craniofacial phenotypes of embryos harvested from the same intercrosses at E13.5 (**Figure 3**; **Table 2**), when facial clefting and hemorrhaging are obvious in embryos with conditional ablation of *Pdgfra* in the NCC lineage^5,6^. As at E10.5, double-heterozygous embryos were significantly overrepresented (19 embryos vs. 8 expected embryos out of 64 total, *χ*^2^ two-tailed *p* = 0.0001) among the harvested embryos, while *Pdgfra^+/fl^;Shp2^fl/fl^;Wnt1-Cre^+/+^* embryos (2 embryos vs. 8 expected embryos, *χ*^2^ two-tailed *p* = 0.023) and *Pdgfra^+/fl^;Shp2^fl/fl^;Wnt1-Cre^+/Tg^* embryos (1 embryo vs. 8 expected embryos, *χ*^2^ two- tailed *p* = 0.0082; **Table 2**) were significantly underrepresented. *Pdgfra^fl/fl^;Shp2^+/fl^;Wnt1-Cre^+/Tg^* and double-homozygous mutant embryos were recovered at Mendelian frequencies at E13.5 (**Table 2**). Double-heterozygous embryos (**Figure 3B,B’**) were indistinguishable from control littermates (**Figure 3A,A’,I,I’**) at E13.5, while *Pdgfra^+/fl^;Shp2^fl/fl^;Wnt1-Cre^+/Tg^* (**Figure 3D,D’**), *Pdgfra^fl/fl^;Shp2^+/fl^;Wnt1-Cre^+/Tg^* (**Figure 3F,F’**) and double-homozygous mutant embryos (**Figure 3H,H’,J,J’**) exhibited variable penetrance of facial blebbing (0%, n = 1; 40%, n= 5; 75%, n = 8), facial hemorrhaging (100%, n = 1; 40%, n= 5; 63%, n = 8), midline clefting (100%, n = 1; 60%, n = 5; 88%, n = 8) and loss of the mandibular region (100%, n = 1; 40%, n = 5; 88%, n = 8) at this timepoint (**Table 2**). In general, the phenotypes of allelic series embryos with homozygous, conditional loss of a particular allele tended to phenocopy single-homozygous mutant embryos of the same allele. For example, *Pdgfra^+/fl^;Shp2^fl/fl^;Wnt1-Cre^+/Tg^* embryos (**Figure 3D,D’**) resembled *Shp2^fl/fl^;Wnt1-Cre^+/Tg^* embryos with loss of most of the upper and lower face^27^, while *Pdgfra^fl/fl^;Shp2^+/fl^;Wnt1-Cre^+/Tg^* embryos (**Figure 3F,F’**) often phenocopied *Pdgfra^fl/fl^;Wnt1-Cre^+/Tg^* embryos with noticeable midline clefting^5,6^. Double-homozygous mutant embryos exhibited a combination, but not a worsening, of the above phenotypes (**Figure 3H,H’,J,J’**). Of note, our findings indicated that SHP2 may negatively regulate PDGFR*α* signaling, as introduction of a *Shp2^fl^* conditional allele partially rescued the craniofacial defects of *Pdgfra* conditional knock-out embryos at E13.5. Specifically, while it has previously been reported that *Pdgfra^fl/fl^;Wnt1-Cre^+/Tg^* embryos at this timepoint exhibit a fully penetrant midline clefting phenotype^5,6^, only 60% of *Pdgfra^fl/fl^;Shp2^+/fl^;Wnt1-Cre^+/Tg^* embryos exhibited midline clefting at E13.5 (**Table 2**).

**Figure 3.**
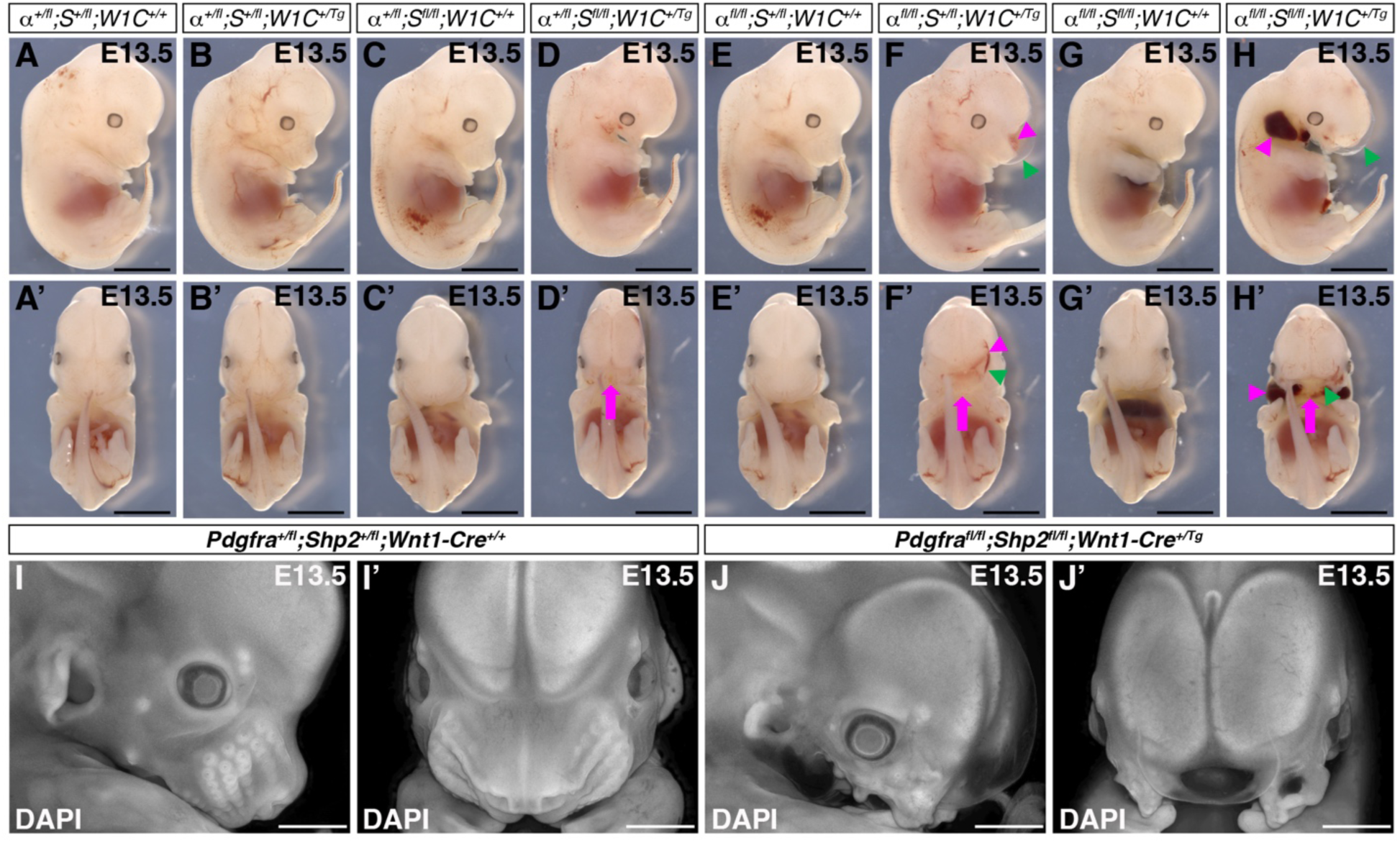
Variably penetrant facial blebbing, facial hemorrhaging, midline clefting and loss of the mandibular region in allelic series mutant embryos at E13.5. (A-J’) Gross morphology of E13.5 embryos resulting from intercrosses of *Pdgfra^fl/fl^;Shp2^fl/fl^* mice with *Pdgfra^+/fl^;Shp2^+/fl^;Wnt1- Cre^+/Tg^* mice viewed laterally (A-J) and frontally (A’-J’) before (A-H’) and after (I-J’) DAPI staining (white). Magenta arrows denote facial clefting; magenta arrowheads denote facial hemorrhaging; green arrowheads denote facial blebbing. The *Pdgfra^+/fl^;Shp2^fl/fl^;Wnt1-Cre^+/Tg^* embryo shown was harvested with a noticeable facial hemorrhage that dissolved during fixation. Scale bars, 1 mm.

**Table 2.** Genotypes and phenotypes of E13.5 embryos from intercrosses of *Pdgfra^fl/fl^;Shp2^fl/fl^* mice with *Pdgfra^+/fl^;Shp2^+/fl^;Wnt1-Cre^+/Tg^* mice.

| Genotype | Expected | Observed | Normal of alive | Dead | Facial bleb | Facial hemorrhage | Midline clefting | Absent mandibular region |
| --- | --- | --- | --- | --- | --- | --- | --- | --- |
| $\alpha^{+/fl};S^{+/fl};W1C^{+/+}$ | 0.125 | 0.188 | 11/11 | 1 | 0/11 | 0/11 | 0/11 | 0/11 |
| $\alpha^{+/fl};S^{+/fl};W1C^{+/Tg}$ | 0.125 | 0.297* | 16/16 | 3 | 0/16 | 0/16 | 0/16 | 0/16 |
| $\alpha^{+/fl};S^{fl/fl};W1C^{+/+}$ | 0.125 | 0.031* | 2/2 | 0 | 0/2 | 0/2 | 0/2 | 0/2 |
| $\alpha^{+/fl};S^{fl/fl};W1C^{+/Tg}$ | 0.125 | 0.016* | 0/1 | 0 | 0/1 | 1/1 | 1/1 | 1/1 |
| $\alpha^{fl/fl};S^{+/fl};W1C^{+/+}$ | 0.125 | 0.078 | 4/5 | 0 | 1/5 | 0/5 | 0/5 | 0/5 |
| $\alpha^{fl/fl};S^{+/fl};W1C^{+/Tg}$ | 0.125 | 0.094 | 2/5 | 1 | 2/5 | 2/5 | 3/5 | 2/5 |
| $\alpha^{fl/fl};S^{fl/fl};W1C^{+/+}$ | 0.125 | 0.125 | 5/5 | 3 | 0/5 | 0/5 | 0/5 | 0/5 |
| $\alpha^{fl/fl};S^{fl/fl};W1C^{+/Tg}$ | 0.125 | 0.172 | 0/8 | 3 | 6/8 | 5/8 | 7/8 | 7/8 |

\* = statistically significant in $\chi^2$ analysis; n = 64 embryos harvested and scored for phenotype

Given the noticeably increased severity of phenotypes across the allelic series of embryos by E13.5, we analyzed cell proliferation and cell death via Ki67 immunofluorescence analysis and TUNEL staining, respectively, at the earlier timepoint of E10.5. Though we observed trends of decreased cell proliferation across the facial processes in multiple allele combinations compared to control embryos, these changes were not statistically significant (**Figure 4A**). Alternatively, we detected increased cell death in each of the experimental genotypes compared to control embryos, with significant changes in the lateral nasal processes (LNP) and maxillary processes (MxP; **Figure 4B-G’**). Specifically, double-heterozygous (*p* = 0.0453; **Figure 4D,D’**) and double-homozygous mutant embryos (*p* = 0.0335; **Figure 4G,G’**) exhibited a higher percentage of TUNEL-positive cells in the LNP compared to control embryos (**Figure 4C,C’**), while *Pdgfra^+/fl^;Shp2^fl/fl^;Wnt1-Cre^+/Tg^* embryos (**Figure 4E,E’**) had the highest percentage of TUNEL-positive cells at each location assayed, including statistically significant increases in both the LNP (*p* = 0.0437) and MxP (*p* < 0.0001) compared to control embryos (**Figure 4B**). These findings identified cell death in the LNP and MxP as a driver of the observed upper jaw phenotypes in embryos with loss of SHP2.

**Figure 4.**
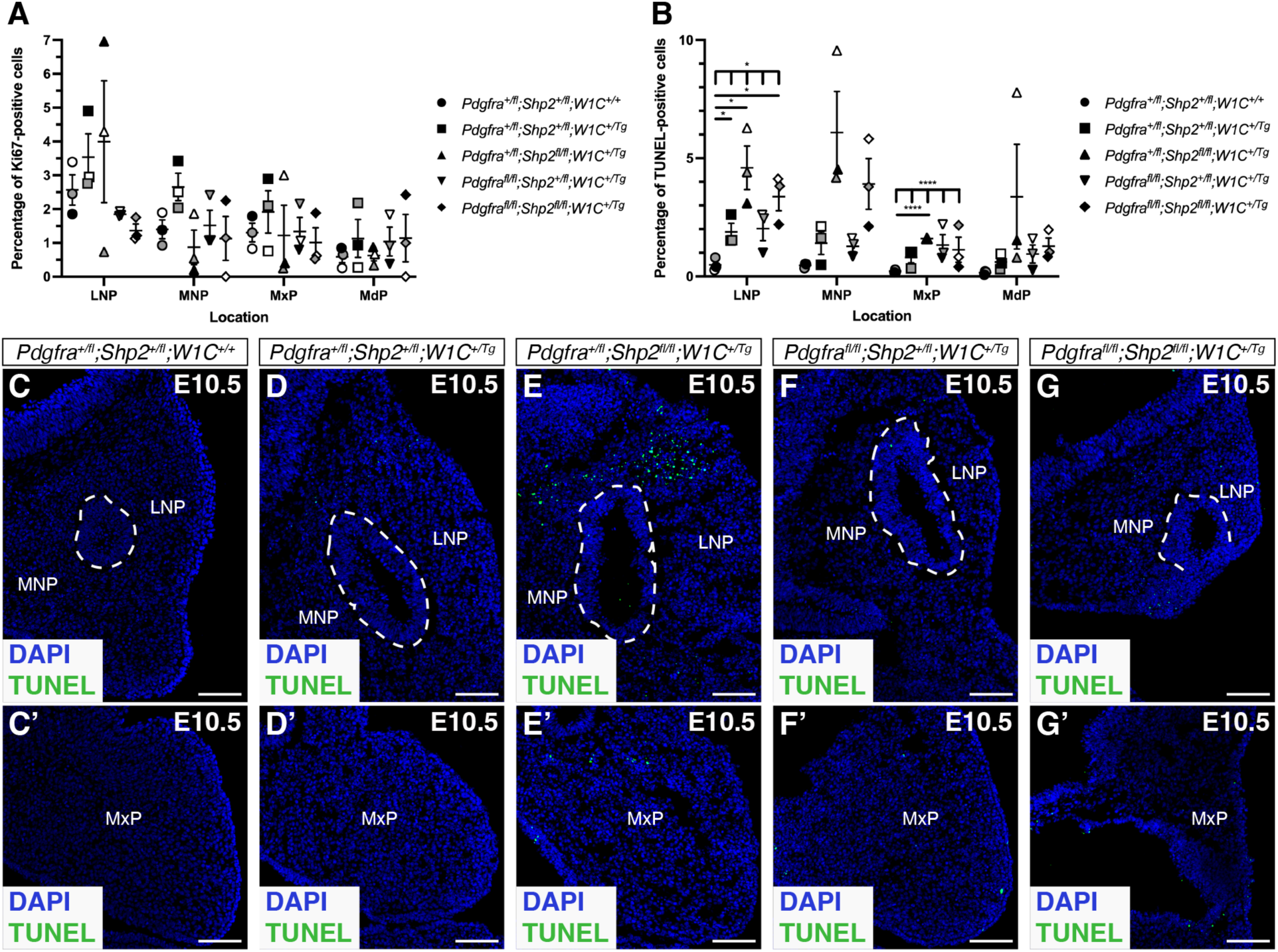
Increased cell death in the lateral nasal and maxillary processes of allelic series mutant embryos at E10.5. (A,B) Scatter dot plots depicting percentages of Ki67-positive (A) and TUNEL-positive (B) cells per embryo in the facial processes across five genotypes at E10.5. Data are presented as mean ± s.e.m. *, p < 0.05; ****, p < 0.0001. Shades correspond to independent experiments. *n* = 3 biological replicates per genotype. LNP, lateral nasal process; MNP, medial nasal process; MxP, maxillary process; MdP, mandibular process. (C-G’) TUNEL staining (green) on sections of medial and lateral nasal processes (C-G) and maxillary processes (C’-G’) across five genotypes at E10.5. Nuclei were stained with DAPI (blue). White dashed lines outline nasal pit epithelium. Scale bars, 100 μm.

We next assessed the craniofacial phenotypes of pups collected from the same intercrosses at postnatal day 0 (P0; **Figure 5**; **Table 3**). At this timepoint, control (19 pups vs. 8 expected pups out of 61 total, *χ*^2^ two-tailed *p* = 0.0001) and double-heterozygous neonates (13 pups vs. 8 expected pups, *χ*^2^ two-tailed *p* = 0.037) were again significantly overrepresented among the harvested progeny, while *Pdgfra^+/fl^;Shp2^fl/fl^;Wnt1-Cre^+/+^* pups were again significantly underrepresented (1 pup vs. 8 expected pups, *χ*^2^ two-tailed *p* = 0.010; **Table 3**). All other genotypes were recovered at Mendelian frequencies (**Table 3**). Double-heterozygous pups (**Figure 5B-B”**) were again indistinguishable from control littermates (**Figure 5A-A”**) at birth. As previously reported for *Pdgfra^fl/fl^;Wnt1-Cre^+/Tg^* mutant skeletons^5,6^, *Pdgfra^fl/fl^;Shp2^+/fl^;Wnt1-Cre^+/Tg^* neonates exhibited bone and cartilage defects, particularly at the midline (**Figure 5F-F”**). Specifically, these pups had fully penetrant clefting of the nasal cartilage, premaxilla, palatal process of the maxilla, and palatal process of the palatine. Further, the nasal cartilage was upturned and the squamosal, caudal process of squamosal, alisphenoid, pterygoid and basisphenoid bones were hypomorphic and misshapen in all *Pdgfra^fl/fl^;Shp2^+/fl^;Wnt1-Cre^+/Tg^* neonates examined (**Figure 5F-F”**). Again, our results indicated that SHP2 may negatively regulate PDGFR*α* signaling, as introduction of a *Shp2^fl^* conditional allele here rescued the nasal cartilage and premaxilla shortening observed in *Pdgfra* conditional knock-out embryos at late gestation^5,6^ (**Figure 5F-F”**).

**Figure 5.**
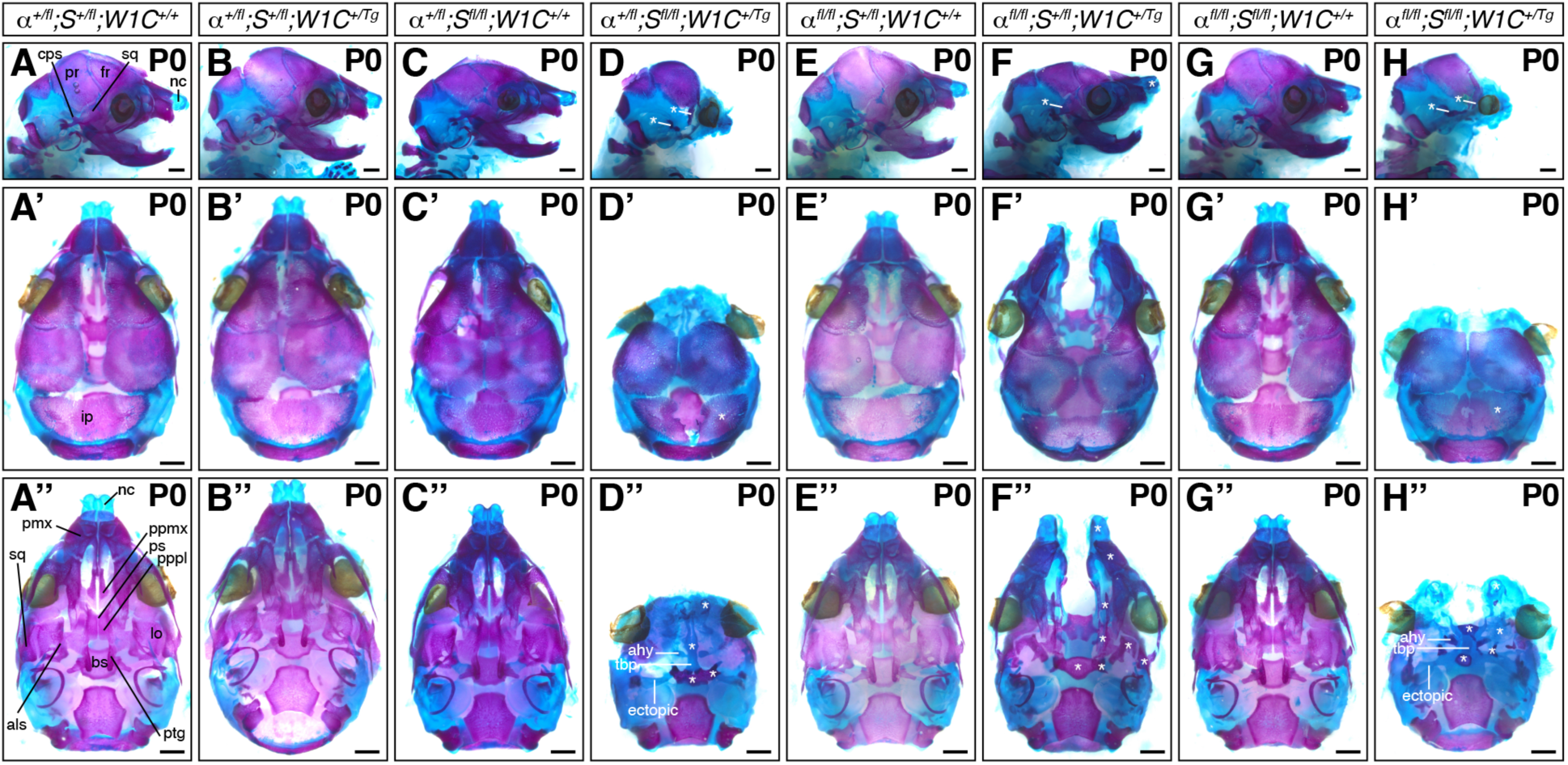
Bone and cartilage phenotypes in allelic series mutant embryos at P0. (A-H’’) Craniofacial skeletal preparations of P0 neonates resulting from intercrosses of *Pdgfra^fl/fl^;Shp2^fl/fl^* mice with *Pdgfra^+/fl^;Shp2^+/fl^;Wnt1-Cre^+/Tg^* mice viewed laterally (A-H), dorsally (A’-H’) and ventrally (A’’-H’’). nc, nasal cartilage; sq, squamosal; fr, frontal; pr, parietal; cps, caudal process of squamosal; ip, interparietal; pmx, premaxilla; ppmx, palatal process of maxilla; ps, presphenoid; pppl, palatal process of palatine; lo, lamina obturans; als, alisphenoid; bs, basisphenoid; ptg, pterygoid; ahy, ala hypochiasmata; tbp, trabecular basal plate. White asterisks indicate abnormal structures compared to those in *Pdgfra^+/fl^;Shp2^+/fl^;Wnt1-Cre^+/+^* control neonates. Scale bars, 1 mm.

**Table 3.** Genotypes and phenotypes of P0 neonates from intercrosses of *Pdgfra^fl/fl^;Shp2^fl/fl^* mice with *Pdgfra^+/fl^;Shp2^+/fl^;Wnt1-Cre^+/Tg^* mice.

| Genotype | Expected | Observed | Normal of alive | Dead | Midline clefting | Absent maxilla | Absent mandible |
| --- | --- | --- | --- | --- | --- | --- | --- |
| $\alpha^{+/fl};S^{+/fl};W1C^{+/+}$ | 0.125 | 0.311* | 19/19 | 0 | 0/19 | 0/19 | 0/19 |
| $\alpha^{+/fl};S^{+/fl};W1C^{+/Tg}$ | 0.125 | 0.213* | 12/12 | 1 | 0/12 | 0/12 | 0/12 |
| $\alpha^{+/fl};S^{fl/fl};W1C^{+/+}$ | 0.125 | 0.016* | 1/1 | 0 | 0/1 | 0/1 | 0/1 |
| $\alpha^{+/fl};S^{fl/fl};W1C^{+/Tg}$ | 0.125 | 0.066 | 0/0 | 4 | 0/0 | 0/0 | 0/0 |
| $\alpha^{fl/fl}; S^{+/fl}; W1C^{+/+}$ | 0.125 | 0.066 | 4/4 | 0 | 0/4 | 0/4 | 0/4 |
| $\alpha^{fl/fl}; S^{+/fl}; W1C^{+/Tg}$ | 0.125 | 0.049 | 0/2 | 1 | 2/2 | 0/2 | 0/2 |
| $\alpha^{fl/fl}; S^{fl/fl}; W1C^{+/+}$ | 0.125 | 0.197 | 11/11 | 1 | 0/11 | 0/11 | 0/11 |
| $\alpha^{fl/fl}; S^{fl/fl}; W1C^{+/Tg}$ | 0.125 | 0.082 | 0/0 | 5 | 0/0 | 0/0 | 0/0 |

\* = statistically significant in $\chi^2$ analysis; n = 61 neonates harvested and scored for phenotype

Importantly, we did not recover any live *Pdgfra^+/fl^;Shp2^fl/fl^;Wnt1-Cre^+/Tg^* nor double-homozygous mutant pups at P0 (**Table 3**). Combined, the progeny ratios that we observed indicate that *Pdgfra^+/fl^;Shp2^fl/fl^;Wnt1-Cre^+/Tg^* embryos die throughout embryogenesis, from before E10.5 to just before birth (**Tables 1-3**). Alternatively, live double-homozygous mutant pups were recovered above or at Mendelian frequencies at E10.5 and E13.5, with the majority of embryos dying during late gestation (**Tables 1-3**). While pups of both genotypes exhibited a complete absence of the upper and lower jaws at P0 (**Figure 5D-D”,H-H”**), there were subtle differences in the bone and cartilage phenotypes. Laterally, all collected pups of both genotypes lacked all neural crest-derived bones anterior and ventral to the parietal bone, with the exception of rudimentary frontal and squamosal bones that tended to be slightly more developed in double-homozygous mutant neonates (**Figure 5H**) than *Pdgfra^+/fl^;Shp2^fl/fl^;Wnt1-Cre^+/Tg^* neonates (**Figure 5D**). Further, the interparietal bone, while somewhat discontinuous in double-homozygous mutant neonates (**Figure 5H’**), was separated into two pieces in most (75%, n = 4) *Pdgfra^+/fl^;Shp2^fl/fl^;Wnt1-Cre^+/Tg^* neonates (**Figure 5D’**). From a ventral perspective, *Pdgfra^+/fl^;Shp2^fl/fl^;Wnt1-Cre^+/Tg^* neonates exhibited nearly adjacent nasal cartilage (**Figure 5D’’**) while all double-homozygous mutant pups had more pronounced clefting of the nasal cartilage (**Figure 5H’’**). Moreover, the trabecular basal plate was evident in all neonates of both genotypes continuous with the presphenoid and ala hypochiasmata, which were variably ossified in *Pdgfra^+/fl^;Shp2^fl/fl^;Wnt1-Cre^+/Tg^* (75%, n = 4; **Figure 5D’’**) and double-homozygous mutant (80%, n = 5; **Figure 5H’’**) neonates. While the basisphenoid and pterygoid bones were hypomorphic in all neonates, they were less developed in double-homozygous mutant pups (**Figure 5H’’**) than *Pdgfra^+/fl^;Shp2^fl/fl^;Wnt1-Cre^+/Tg^* pups (**Figure 5D’’**). Interestingly, ectopic cartilage struts attached to the trabecular basal plate extended posterolaterally towards the middle ear in all pups of both genotypes (**Figure 5D’’,H’’**). Finally, double-homozygous mutant pups displayed incompletely penetrant unossified lamina obturans (80%, n = 5; **Figure 5H’’**). The cartilage elements in both *Pdgfra^+/fl^;Shp2^fl/fl^;Wnt1-Cre^+/Tg^* and double-homozygous mutant neonates were more reminiscent of the late embryonic chondrocranium than that typically observed in neonatal pups^39^, consistent with the fact that these embryos survived to late embryogenesis but were not alive at birth. Further, the presence of unossified elements indicated that bone matrix was not appropriately deposited and/or mineralized.

To explore a possible molecular mechanism underlying the above phenotypes, we examined phospho-PDGFR and phospho-Erk1/2 levels via western blotting of whole-cell lysates derived from the facial processes of E10.5 allelic series embryos. Relative phospho-PDGFR and phospho-Erk1/2 levels were determined by normalizing to total PDGFR*α*and total Erk1/2 levels, respectively. We observed relatively equal levels of phospho-PDGFR across *Pdgfra^+/fl^;Wnt1-Cre^+/Tg^* (1.7 ± 0.29 relative induction (R.I.); mean ± s.e.m.), *Pdgfra^+/fl^;Shp2^+/fl^;Wnt1-Cre^+/+^* (1.0 ± 0.0 R.I.) and double-heterozygous (1.3 ± 0.36 R.I.) samples, with a noticeable increase in *Pdgfra^+/fl^;Shp2^fl/fl^;Wnt1-Cre^+/Tg^* (4.0 ± 1.9 R.I.) lysates (**Figure 6A,B**). Finally, we explored phospho-Erk1/2 levels in the same lysates, revealing progressively increasing relative induction values across *Pdgfra^+/fl^;Wnt1-Cre^+/Tg^* (1.1 ± 0.32 R.I.), double-heterozygous (1.4 ± 0.28 R.I.) and *Pdgfra^+/fl^;Shp2^fl/fl^;Wnt1-Cre^+/Tg^* (2.0 ± 1.2 R.I.) samples (**Figure 6C,D**). Together, these findings demonstrated that loss of SHP2 leads to greater phosphorylation of PDGFR*α* and increased phospho-Erk1/2 levels.

**Figure 6.**
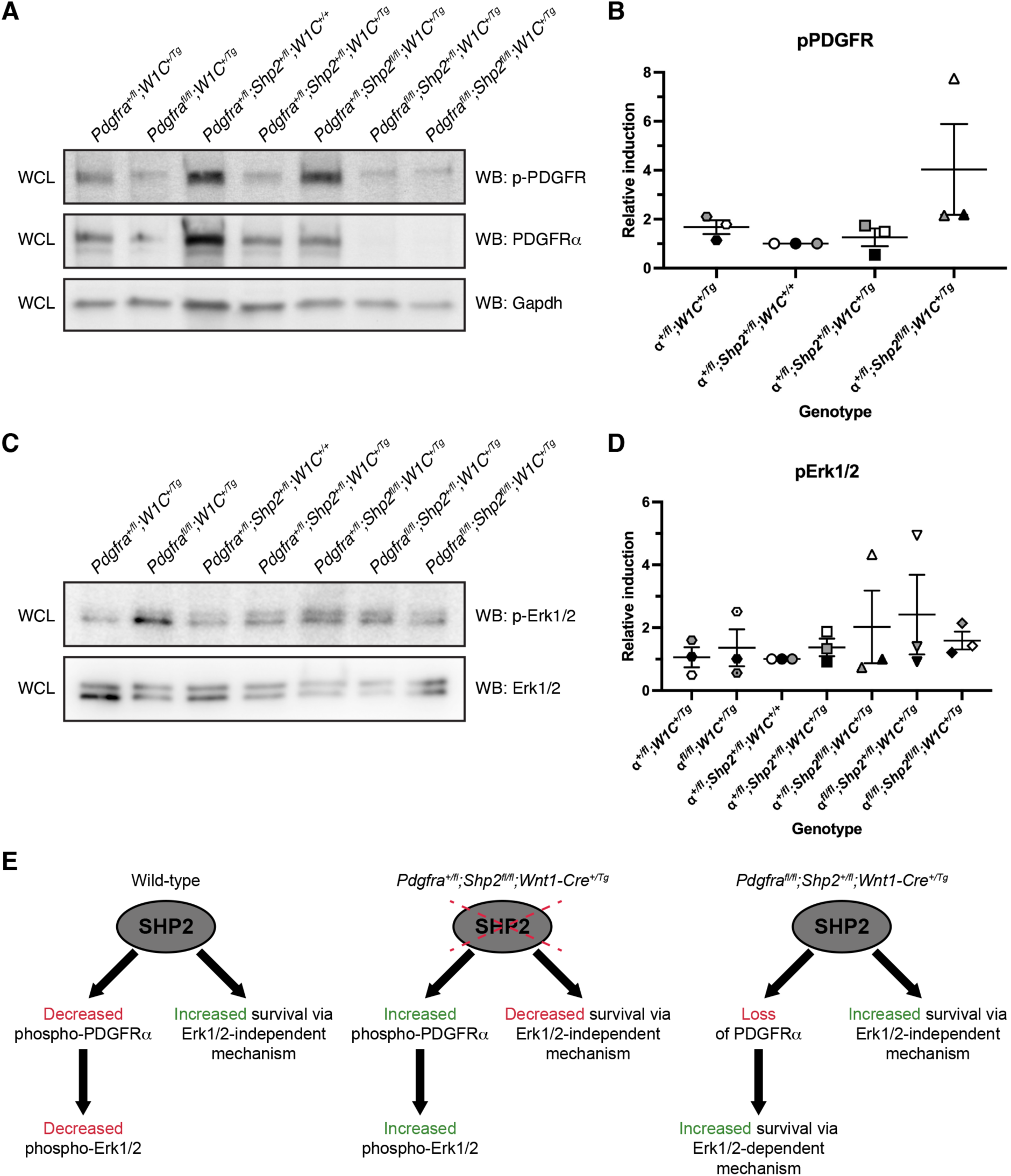
Loss of SHP2 leads to greater phosphorylation of PDGFR*α*and increased phospho- Erk1/2 levels. (A) Western blotting of whole-cell lysates (WCL) with anti-phospho(p)-PDGFR, anti-PDGFR*α* and anti-Gapdh antibodies. (B) Scatter dot plot depicting quantification of band intensities from three independent experiments as in A. Data are presented as mean ± s.e.m. Shades correspond to independent experiments. *n* = 3 biological replicates per condition. (C) Western blotting of WCL with anti-phospho(p)-Erk1/2 and Erk1/2 antibodies. (D) Scatter dot plot depicting quantification of band intensities from three independent experiments as in C. Data are presented as mean ± s.e.m. Shades correspond to independent experiments. *n* = 3 biological replicates per condition. (E) Proposed model in which SHP2 binds and dephosphorylates PDGFR*α* while simultaneously increasing survival through an Erk1/2- independent mechanism.

## Discussion

Our results demonstrated that the ablations examined here are largely additive in nature, as double-homozygous mutant embryos exhibited a combination, but not an improvement or worsening, of the previously reported phenotypes observed upon conditional loss of PDGFR*α* or SHP2 in the mouse NCC lineage. This finding underscores PDGFR*α*-independent functions of SHP2 in this context and is perhaps not surprising given the demonstrated dependencies of PDGFR*α* on PI3K/Akt signaling^7,11–13^ and of SHP2 on Erk1/2 signaling^27,31,32^ in the mouse NCC lineage, as well as the large number of RTKs active in this cell type^40^. However, we provided several lines of evidence that SHP2 may negatively regulate PDGFR*α* signaling. First, while *Pdgfra^fl/fl^;Shp2^+/fl^;Wnt1-Cre^+/+^* embryos with no Cre expression were recovered below expected levels at all timepoints assayed, and significantly so at E10.5, introduction of a *Cre* allele in *Pdgfra^fl/fl^;Shp2^+/fl^;Wnt1-Cre^+/Tg^* embryos improved recovery during embryogenesis. The rate of recovery of the former genotype is not unexpected, as we and others have previously reported that the *Pdgfra^fl^* allele is hypomorphic^5,10,41^. Moreover, double-homozygous mutant embryos were recovered at increased frequencies compared to both *Pdgfra^fl/fl^;Shp2^+/fl^;Wnt1-Cre^+/+^* embryos and *Pdgfra^fl/fl^;Shp2^+/fl^;Wnt1-Cre^+/Tg^* embryos at E10.5 and E13.5. Second, *Pdgfra^fl/fl^;Shp2^+/fl^;Wnt1-Cre^+/Tg^* embryos exhibited decreased penetrance of midline facial clefting at E13.5 as well as rescue of nasal cartilage and premaxilla shortening at P0 compared to *Pdgfra^fl/fl^;Wnt1-Cre^+/Tg^* embryos at these same timepoints. It is unlikely that these effects are explained by subtle differences in genetic background, as *Pdgfra^GFP/GFP^* embryos in our colony on pure 129S4 or C57BL/6 backgrounds exhibit fully penetrant facial clefting. Third, phospho-PDGFR and phospho-Erk1/2 levels were modestly increased in E10.5 *Pdgfra^+/fl^;Shp2^fl/fl^;Wnt1- Cre^+/Tg^* lysates relative to double-heterozygous and *Pdgfra^+/fl^;Wnt1-Cre^+/Tg^* samples.

Alternatively, results from this study also indicated that SHP2 may positively regulate PDGFR*α* signaling. Though the rate of recovery was improved upon addition of conditional *Shp2^fl^* alleles in the absence of PDGFR*α*, phenotypes of the resulting progeny worsened, such that *Pdgfra^fl/fl^;Shp2^+/fl^;Wnt1-Cre^+/+^* embryos had fewer defects than *Pdgfra^fl/fl^;Shp2^+/fl^;Wnt1- Cre^+/Tg^* embryos, which in turn had less severe phenotypes than double-homozygous mutant embryos. Further, *Pdgfra^+/fl^;Shp2^fl/fl^;Wnt1-Cre^+/Tg^* embryos and neonates were recovered at decreased frequencies and had much more severe craniofacial phenotypes compared to double-heterozygous mice at all timepoints examined. Of note, the fact that *Pdgfra^+/fl^;Shp2^fl/fl^;Wnt1-Cre^+/+^* embryos were underrepresented was unexpected, as there are no reports in the literature that the *Shp2^fl^* allele is hypomorphic and *Shp2^fl/fl^* mice in our colony, which are maintained through homozygous intercrosses, give birth to litters of expected sizes (average litter size of 5.3 pups at 5-10 days after birth compared to an average of 5.4 pups for wild-type 129S4 litters, p = 0.70).

These seemingly contradictory results can be explained by a model in which SHP2 binds and dephosphorylates PDGFR*α*while simultaneously increasing survival through an Erk1/2-independent mechanism (**Figure 6E**). SHP2 cannot dephosphorylate PDGFR*α* in a situation in which the receptor has been ablated. Instead, in the case of *Pdgfra^fl/fl^;Shp2^+/fl^;Wnt1- Cre^+/Tg^* mice it is likely that loss of PDGFR*α* frees up SHP2 and other signaling molecules that bind the receptor to contribute to cell survival and/or skeletal development via Erk1/2-dependent and -independent mechanisms. Consistent with this idea, phospho-Erk1/2 levels were the highest in lysates from *Pdgfra^fl/fl^;Shp2^+/fl^;Wnt1-Cre^+/Tg^* embryos compared to all other assayed genotypes. Interestingly, *Pdgfra^+/fl^;Shp2^fl/fl^;Wnt1-Cre^+/Tg^* mice had the most severe phenotypes among all genotypes analyzed across several metrics. These included the rate of recovery at E10.5-P0, the percentage of TUNEL-positive cells in the facial processes at E10.5 and phenotypes of skeletal elements such as the frontal, squamosal and interparietal bones at P0.

Interestingly, however, levels of Erk1/2 phosphorylation were essentially equal in *Pdgfra^+/fl^;Shp2^fl/fl^;Wnt1-Cre^+/Tg^* and double-homozygous mutant lysates at E10.5. Here, it is likely that loss of SHP2 in the presence of PDGFR*α* results in modest increases in phospho-PDGFR and phospho-Erk1/2 levels that are not sufficient to offset Erk1/2-independent functions of SHP2 in preventing cell death. Moreover, it is possible that sustained phosphorylation of PDGFR*α* may negatively contribute to cell survival and skeletal development through altered interaction dynamics with additional signaling molecules. These interactions would not be possible in double-homozygous mutant embryos, potentially explaining the improved recovery of this genotype compared to *Pdgfra^+/fl^;Shp2^fl/fl^;Wnt1-Cre^+/Tg^* mice. Further studies would be required to determine the signaling molecule(s) underlying these mechanisms.

Previous studies have revealed that both PDGFR*α* and SHP2 play a role in the differentiation of NCC-derived cells into bone, and that this process requires Erk1/2. Stimulation of cultured primary mouse embryonic palatal mesenchyme cells with PDGF-AA ligand led to transcriptional changes associated with skeletal differentiation and expression of the osteoblast marker alkaline phosphatase^8^. Further, E17.5 embryos with NCC-specific ablation of SHP2 exhibited loss of expression of the osteoblast marker osteopontin in the frontal bones^27^. Reduced osteopontin expression was also detected in the frontal bones of P3 neonates with a NS1-associated *Ptpn11* variant active in the NCC lineage^32^. Given the fact that the above SHP2 models have altered phospho-Erk1/2 levels in structures ranging from the mid-gestation pharyngeal arches^27^ to the neonatal frontal bones^32^ and that osteopontin expression levels were rescued in the latter model upon treatment with the MEK1/2 inhibitor U0126^32^, these findings indicate that precise phospho-Erk1/2 levels are required for proper differentiation of mouse cranial NCC-derived mesenchyme into bone. Consistent with this hypothesis, NCC-specific ablation of Erk2 leads to decreased expression of the osteogenic progenitor marker Sp7 and the differentiated osteoblast marker *Col1a1* in the mandible past mid-gestation, resulting in micrognathia and mandibular asymmetry^42^.

Importantly, an Erk1/2-independent mechanism of SHP2 activity that contributes to cell survival has previously been identified in zebrafish neural crest^43^. Specifically, SHP2 was shown to inhibit a p53-dependent cell death pathway in this context, independent of both SHP2 catalytic activity and Erk1/2 activation. This role is separate from a phosphatase- and Erk1/2- dependent function of SHP2 that contributes to NCC specification and migration^43^. Interestingly, we have previously shown that PDGFR*α* signaling regulates survival in mouse skeletal development through p53-dependent intracellular pathways^7^. Whether SHP2 and PDGFR*α* signaling converge on a shared p53 node in the neural crest and their derivatives remains to be determined, though such an interaction is unlikely given the additive nature of our phenotypic results. However, our findings indicate that an Erk1/2-independent mechanism of survival mediated by SHP2 may be conserved in the mouse neural crest lineage.

## Experimental Procedures

### Transient transfection

HEK 293T/17 cells (American Type Culture Collection) were cultured in medium [Dulbecco’s modified Eagle’s medium (Gibco, Thermo Fisher Scientific, Waltham, MA) supplemented with 50 U/mL penicillin (Gibco), 50 μg/mL streptomycin (Gibco) and 2 mM L-glutamine (Gibco)] containing 10% FBS (Hyclone Laboratories Inc., Logan, UT) at 37°C in 5% CO_2_. Cells at 70% confluence were transfected with combinations of pDEST-PDGFR-V1^34^ and pDEST-PDGFR-V2 constructs^34^ (5 μg each) or pCAG:myr-Venus^37^ (#32602; Addgene; 5 μg) using Lipofectamine LTX (Thermo Fisher Scientific).

### Immunoprecipitation and western blotting

To induce PDGFR signaling 24 h after transient transfection, HEK 293T/17 cells were serum starved for 24 h in medium containing 0.1% FBS and stimulated with 10 ng/mL PDGF- AA (PDGFR*α*/*α*homodimers) or PDGF-BB (PDGFR*α*/*β*heterodimers and PDGFR*β*/*β*homodimers) ligand (R&D Systems, Minneapolis, MN) for 15 min. Protein lysates were generated by resuspending cells in GFP-Trap lysis buffer [20 mM Tris-HCl pH 7.5, 150 mM NaCl, 1 mM EDTA, 0.5% Nonidet P-40, 1 mM PMSF and 1x complete Mini protease inhibitor cocktail (Roche, MilliporeSigma, Burlington, MA)] and collecting cleared lysates by centrifugation at 13,400 g at 4°C for 20 min. For immunoprecipitations, cell lysates (500 μg) were incubated with GFP-Trap agarose beads (Bulldog Bio, Inc., Portsmouth, NH) for 1 hr at 4°C. Beads were washed three times with ice-cold GFP-Trap wash/dilution buffer [10 mM Tris-HCl pH 7.5, 150 mM NaCl, 0.5 mM EDTA] and the precipitated proteins were eluted with Laemmli buffer containing 10% *β*-mercaptoethanol, heated for 10 min at 100°C and separated by SDS-PAGE. Facial processes (including the MNP, LNP, MxP and MdP) were dissected on ice from E10.5 embryos^44^, snap frozen in 100% ethanol on dry ice and stored at -80°C. Protein lysates were generated by resuspending tissues from 1-2 embryos in ice-cold NP-40 lysis buffer [20 mM Tris-HCl (pH 8), 150 mM NaCl, 10% glycerol, 1% Nonidet P-40, 2 mM EDTA, 1x complete Mini protease inhibitor cocktail (Roche, MilliporeSigma, Burlington, MA), 1 mM PMSF, 10 mM NaF, 1 mM Na_3_VO_4_ and 25 mM *β*-glycerophosphate] and collecting cleared lysates by centrifugation at 13,4000 g at 4°C for 20 min. Laemmli buffer containing 10% *β*- mercaptoethanol was added to the lysates, which were heated for 5 min at 100°C. Proteins were subsequently separated by SDS-PAGE. Western blot analysis was performed according to standard protocols using horseradish peroxidase-conjugated secondary antibodies. The following antibodies were used for western blotting: SHP2 (1:2,000; AF1894; R&D Systems); PDGFR*α* (1:200; C-20; sc-338; Santa Cruz Biotechnology, Inc., Dallas, TX); PDGFR*β*(1:200; 958; sc-432; Santa Cruz Biotechnology, Inc.); phospho-PDGFR*α* (Tyr849)/PDGFR*β* (Tyr857) (1:1,000; C43E9; 3170; Cell Signaling Technology, Inc., Danvers, MA); PDGFR*α* (1:200; AF1062; R&D Systems); phospho-Erk1/2 (1:1,000; Thr202/Tyr204; 9101; Cell Signaling Technology, Inc.); Erk1/2 (1:1,000; 9102; Cell Signaling Technology, Inc.); horseradish peroxidase-conjugated donkey anti-goat IgG (1:5,000; sc-2033; Santa Cruz Biotechnology, Inc.); horseradish peroxidase-conjugated goat anti-rabbit IgG (1:20,000; 111035003; Jackson ImmunoResearch Inc., West Grove, PA). Quantifications of signal intensity were performed with ImageJ software (version 1.53k, National Institutes of Health, Bethesda, MD). Relative SHP2 binding was determined by normalizing GFP-Trap immunoprecipitated SHP2 levels to total SHP2 levels. Relative phospho-PDGFR and phospho-Erk1/2 levels were determined by normalizing to total PDGFR*α* and total Erk1/2 levels, respectively. Statistical analyses were performed with Prism 9 (GraphPad Software, Inc., La Jolla, CA) using a two-tailed, unpaired *t*- test with Welch’s correction. Immunoprecipitation and western blotting experiments were performed across three independent experiments.

### Mouse strains and husbandry

All animal experimentation was approved by the Institutional Animal Care and Use Committee of the University of Colorado Anschutz Medical Campus. *Pdgfra^tm8Sor^* mice^5^, referred to in the text as *Pdgfra^fl^*; *Ptpn11^tm3Bgn^* mice^45^, referred to in the text as *Shp2^fl^*; and *H2az2^Tg^*(Wnt1–cre)*^11Rth^* mice^4^, referred to in the text as *Wnt1-Cre^Tg^*, were used in this study and housed at a sub-thermoneutral temperature of 21-23°C with *ad libitum* food and water. Progeny were obtained from intercrosses of *Pdgfra^fl/fl^;Shp2^fl/fl^* male mice on a 75% 129S4 genetic background with *Pdgfra^+/fl^;Shp2^+/fl^;Wnt1-Cre^+/Tg^* female mice on a 87.5% 129S4 genetic background. Mice were euthanized by inhalation of carbon dioxide from compressed gas. Cervical dislocation was used as a secondary verification of death. Statistical analyses of Mendelian inheritance were performed with the GraphPad QuickCalcs data analysis resource (GraphPad Software, Inc.) using a *χ*^2^ test.

### Morphological analysis

Embryos were dissected at multiple timepoints (day of plug considered 0.5 days) in 1x phosphate buffered saline (PBS) and fixed overnight at 4°C in 4% paraformaldehyde (PFA) in PBS. Embryos were photographed using an Axiocam 105 color digital camera (Carl Zeiss, Inc., Thornwood, NY) fitted onto a Stemi 508 stereo microscope (Carl Zeiss, Inc.). Distances between nasal pits and heights of heads were measured using Photoshop software (Adobe, San Jose, CA). The normalized distance between nasal pits was calculated by dividing the distance between nasal pits by the height of the head from the anterior surface of the forebrain to the posterior surface of the nasal processes. Statistical analyses were performed with Prism 9 (GraphPad Software, Inc.) using a two-tailed, unpaired *t-*test with Welch’s correction and Welch and Brown-Forsythe ANOVA tests.

### Ki67 immunofluorescence analysis

Embryos were fixed in 4% PFA in PBS and infiltrated with 30% sucrose in PBS before being mounted in O.C.T. compound (Sakura Finetek United States Inc., Torrance, CA). Sections (8 μm) were deposited on glass slides. Sections were fixed in 4% PFA in PBS with 0.1% Triton X-100 for 10 min and washed in PBS with 0.1% Triton X-100. Sections were blocked for 1 h in 5% normal donkey serum (Jackson ImmunoResearch Inc., West Grove, PA) in PBS and incubated overnight at 4°C in anti-Ki67 primary antibody (1:300; PA1-38032; Invitrogen, Carlsbad, CA) in 1% normal donkey serum in PBS. After washing in PBS, sections were incubated in Alexa Fluor 488-conjugated donkey anti-rabbit secondary antibody (1:1,000; A21206 Invitrogen) diluted in 1% normal donkey serum in PBS with 2 μg/mL 4’,6-diamidino-2-phenylindole (DAPI; Sigma-Aldrich Corp., St. Louis, MO) for 1 h. Sections were mounted in Aqua Poly/Mount mounting medium (Polysciences, Inc., Warrington, PA) and photographed using an Axiocam 506 mono digital camera (Carl Zeiss, Inc.) fitted onto an Axio Observer 7 fluorescence microscope (Carl Zeiss, Inc.). All positive signals were confirmed by DAPI staining. The percentage of Ki67-positive cells was determined in three embryos per genotype. Statistical analyses were performed with Prism 9 (GraphPad Software, Inc.) using a two-tailed, unpaired *t-* test with Welch’s correction and Welch and Brown-Forsythe ANOVA tests.

### TUNEL assay

Sections (8 μm) of PFA-fixed, sucrose-infiltrated, O.C.T.-mounted embryos were deposited on glass slides. Apoptotic cells were identified using the *In Situ* Cell Death Detection Kit, Fluorescein (Sigma-Aldrich Corp.) according to the manufacturer’s instructions for the treatment of cryopreserved tissues sections. Sections were mounted in VECTASHIELD Mounting Medium for Fluorescence with DAPI (Vector Laboratories, Burlingame, CA) and photographed using an Axiocam 506 mono digital camera (Carl Zeiss, Inc.) fitted onto an Axio Observer 7 fluorescence microscope (Carl Zeiss, Inc.). All positive signals were confirmed by DAPI staining. The percentage of terminal deoxynucleotidyl transferase-mediated dUTP nick end labeling (TUNEL)-positive cells was determined in three embryos per genotype. Statistical analyses were performed with Prism 9 (GraphPad Software, Inc.) using a two-tailed, unpaired *t-* test with Welch’s correction and Welch and Brown-Forsythe ANOVA tests.

### Skeletal preparations

Neonatal pups were skinned, eviscerated, fixed in 95% ethanol and stained in 0.015% Alcian blue, 0.005% Alizarin red and 5% glacial acetic acid in 70% ethanol at 37°C. Pups were then cleared in 1% KOH and transferred to solutions of decreasing KOH concentration and increasing glycerol concentration. Skeletons were photographed using an Axiocam 105 color digital camera (Carl Zeiss, Inc.) fitted onto a Stemi 508 stereo microscope (Carl Zeiss, Inc.).

## Acknowledgments

We are grateful to Erin Binne for technical assistance. The pCAG:myr-Venus plasmid was a gift from Dr. Rytis Prekeris at the University of Colorado Anschutz Medical Campus. *Shp2^fl^* mice were a gift from Dr. Benjamin Neel at New York University Langone Health. We are grateful to Dr. David Clouthier for discussions on the bone and cartilage phenotypes of allelic series neonates. We thank members of the Fantauzzo laboratory for their helpful discussions and critical comments on the manuscript. This work was supported by the National Institutes of Health/National Institute of Dental and Craniofacial Research (R01DE027689 to K.A.F. and K02DE028572 to K.A.F.).

## Author Contributions

Conceptualization: K.A.F.; Methodology: K.A.F.; Formal analysis: D.F., J.J., E.P.B., K.A.F.; Investigation: D.F., J.J., E.P.B., K.A.F.; Writing – Original Draft: K.A.F.; Writing – Reviewing and Editing: D.F., J.J., E.P.B.; Visualization: D.F., J.J., K.A.F.; Supervision: K.A.F.; Funding acquisition: K.A.F.

## Data Availability Statement

All relevant data are provided within the manuscript.

**Grant Sponsor**: National Institute of Dental and Craniofacial Research

**Grant Number**: R01DE027689, K02DE028572

## Conflict of Interest Disclosure

The authors declare no conflict of interest, financial or otherwise.

## Ethics Approval Statement

All animal experimentation was approved by the Institutional Animal Care and Use Committee of the University of Colorado Anschutz Medical Campus.

## Notes

### Competing Interest Statement

The authors have declared no competing interest.

